# More insights into the dynamic assembly and rotation of the archaellum motor machinery

**DOI:** 10.64898/2026.10.29.762886

**Authors:** Marie Joest, Frederic Wüllner, Julia Schimpf, Leonie Vollmar, Elisa Steyer, Rashmi Kumariya, Thorsten Hugel, Sonja-Verena Albers

## Abstract

Archaea use the archaellum for swimming motility. Rotation of the archaellum is driven by the ATPase ArlI and the membrane platform protein ArlJ. Structural studies suggest that conformational changes in ArlI power the rotation of ArlJ. However, how ATP hydrolysis is converted into rotation remains poorly understood. Using single-molecule total internal reflection fluorescence (TIRF) microscopy, we show that dimeric ArlJ forms a complex with ArlI. Combining conservation analysis, AlphaFold3 predictions and co-purifications, we identified conserved ArlJ residues required for interaction with ArlI, including a structurally rigid loop and the critical residue R341 at the ArlI interface. Motility assays in *Haloferax volcanii* confirmed these findings. Together, these findings support a model in which a dimeric ArlJ is interacting with ArlI via conserved residues to generate torque.

## Introduction

The archaellum is a specialised cell surface structure found in archaea that enables swimming motility and environmental responsiveness [1]. It consists of a filament and a motor complex that powers both the assembly and the rotation of the filament. The motor complex is the membrane protein ArlJ and the ATPase ArlI. ArlJ contains 7-9 transmembrane helices and is thought to function as the platform protein, anchoring the filament, a role reminiscent of Type IV pilus (T4P) platform proteins such as PilC, suggesting evolutionary conservation across these systems [2,3]. Cytosolic loops of ArlJ are enriched in positively charged residues, which are proposed to serve as interaction sites for ArlI [4]. Functionally, ArlJ is essential for both assembly and function of the archaellum [5–8].

ArlI, in contrast, is a soluble ATPase and the primary energy-converting component of the motor. It assembles into a hexameric ring and is structurally divided into two domains: a conserved C-terminal (CTD) ATPase domain and a more variable N-terminal domain (NTD), connected by a flexible linker [9–11]. The CTD harbours canonical Walker A and B motifs for ATP binding and hydrolysis. The NTD is implicated in protein-protein interactions, particularly with ArlJ, and includes a flexible three-helix bundle that appears critical for membrane localization and interaction, as mutations in this region disrupt both archaellum assembly and motility [9].

Torque generation is driven by ATP hydrolysis within ArlI [12]. Structural studies suggest a cyclic process centred on conformational changes in the hexameric ATPase. Upon ATP binding, an arginine clamp forms, stabilising the ArlI in a closed conformation. Hydrolysis then triggers electrostatic repulsion and transition to an open state, concurrent with rotation of the NTD. Neighbouring subunits facilitate phosphate release, producing a coordinated conformational wave across the hexamer. These events generate vertical displacement of 8-10 angstroms in the protein structure, likely coupled to both filament assembly and rotational force generation [9].

Two main models have been proposed for filament rotation: In the first rotor model, ArlI remains static while its subunits undergo sequential ATP hydrolysis, cycling between open and closed conformations. In the open state, two opposing ArlI monomers each engage one ArlJ monomer. Upon ATP hydrolysis, a conformational change in ArlI transfers these interactions to the next pair of open-state monomers, producing a stepwise rotation, where each ATP hydrolysis induces a 60° rotation of ArlJ, consistent with the sixfold symmetry of ArlI hexamer [13,14]. In the second model, ArlJ and ArlI are tightly coupled and rotate together as a unit. Here, conformational changes upon ATP hydrolysis cause the CTD of opposing ArlI monomers to push outwards against surrounding stator proteins like ArlCDE or ArlX, generating torque that rotates ArlJ and ArlI as a unit, functioning analogously to a bearing system [14].

In both models, ATP hydrolysis is the sole driving force, distinguishing the archaellum from the proton motif force driven bacterial flagellum. Experimentally measured torque values of approximately 160 pN×nm correspond to an energy consumption of 10-18 J per full rotation. This is consistent with the hydrolysis of at least 12 ATP molecules per rotation, in line with a hexameric ATPase powering the motor complex [15,16].

In summary, the rotation of the archaellum is driven by an ATP-powered motor, with the interaction between ArlJ and ArlI at its heart: ArlI converts chemical energy into mechanical motion through coordinated conformational changes, ultimately driving rotation of the filament. Nevertheless, the precise molecular mechanism by which torque is generated remains unresolved. Using single-molecule TIRF microscopy, we show that ArlJ interacts as a dimer with ArlI. Combined with conservation analysis, AlphaFold3 predictions, co-purification, and motility assays in *Haloferax volcanii*, we identified a structurally rigid proximal loop in ArlJ and a direct ArlI interface, with the conserved residue R341 being the most critical residue. Based on this data, we propose a model in which R341 and neighbouring residues of ArlJ engage with ArlI to generate tangential force allowing rotation of the archaellum motor complex.

## Results

### The membrane platform protein ArlJ forms a dimer

Although T4P have been studied extensively in recent years, the oligomeric state of the membrane platform protein has remained unresolved, with only limited biochemical evidence available [17]. To determine the oligomeric state of ArlJ, we established a dedicated single-molecule total internal reflection fluorescence (smTIRF) microscopy assay. Individually purified ArlJ from *Archaeoglobus fulgidus* and ArlI from *Pyrococcus furiosus* were used for the measurements. Notably, the ATPase ArlI from *A. fulgidus* was predominantly monomeric and unstable when heterologously expressed and purified from *E. coli*. Because *A. fulgidus* ArlJ was the only membrane platform protein that could be reliably expressed and purified, we employed ArlI from a different euryarchaeon for the single-molecule TIRF experiments. ArlI from *P. furiosus* was purified as higher-order oligomeric species as shown by size-exclusion chromatography, although we did not observe a clear hexameric assembly and degradation was visible (Supplementary Figure 1 C, D). However, *P. furiosus* ArlI was the only euryarchaeal ArlI we were able to express and purify under compatible buffer and temperature conditions used for ArlJ. For single-molecule measurements, ArlI was immobilized within a microfluidic flow chamber, which was subsequently flushed with fluorophore-labelled ArlJ to permit complex formation (Figure 1 A). The oligomeric state of ArlJ was then inferred by counting photobleaching steps for individual fluorophore-labelled molecules.

**Figure 1.**
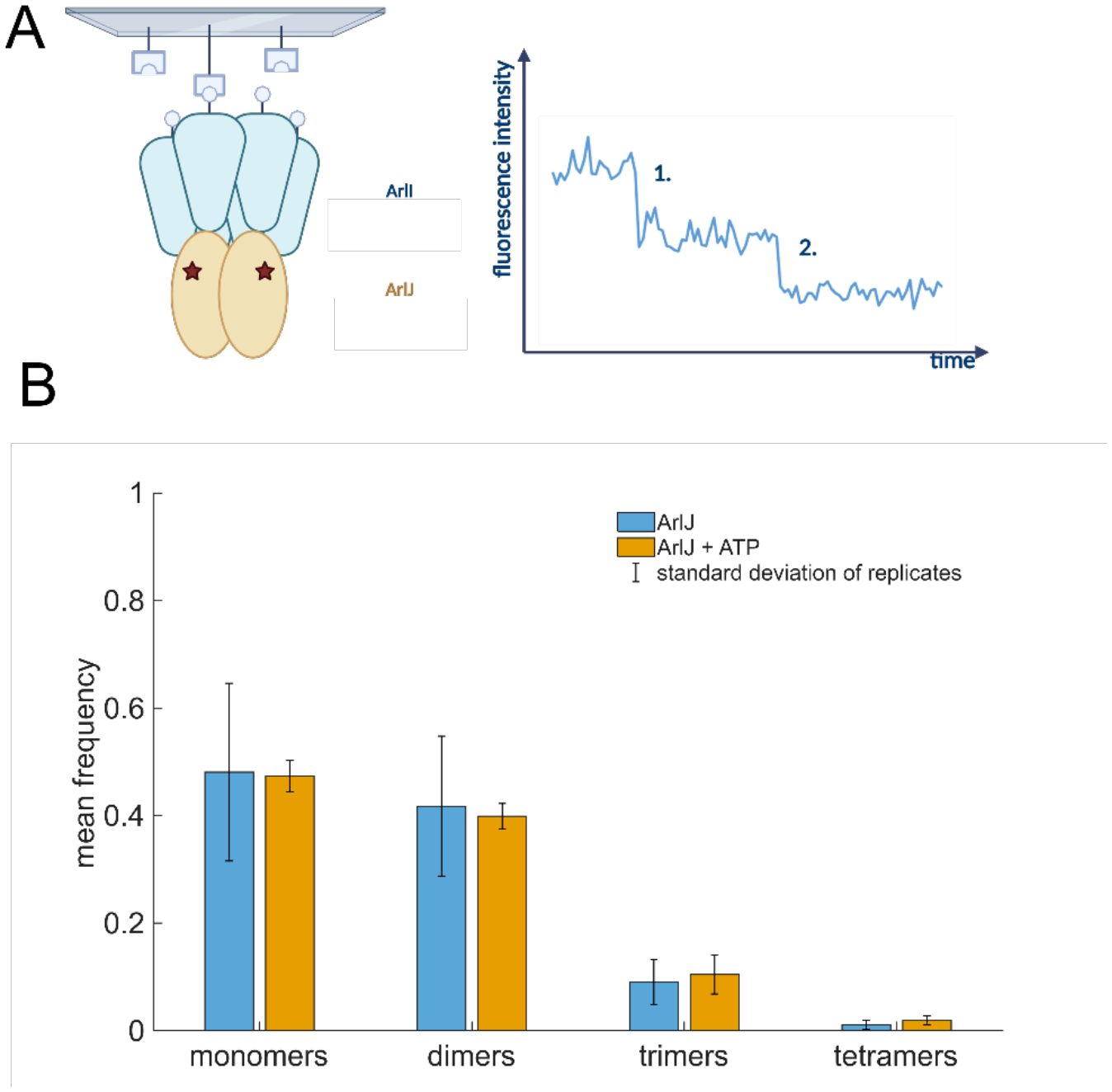
Single-molecule measurements of ArlJ. **A.** Scheme of the experimental set up for the smTIRF measurements. ArlI is immobilized in a microfluidic flow chamber via Biotin-NeutrAvidin. Labelled ArlJ is flushed into the chamber, allowing for complex formation. The chamber is excited with a laser in TIR geometry and bleaching steps are counted over time. **B**. Mean frequency of ArlJ oligomers from bleaching step experiments, measured in the absence (blue) and presence of 1 mM ATP and 5 mM MgCl_2_ (orange). The number of bleaching steps is corrected for the degree of labelling (DOL) of 66 % (see Methods and Materials). For each condition, three independent measurements of biological replicates were performed, in total 1967 traces for the measurements in absence of ATP and 2337 traces for the measurements including ATP were quantified. Figure 1A was created in BioRender. Albers, S. (2027) https://BioRender.com/xnmb9p6.

These measurements showed a predominantly existence of ArlJ as a mixture of monomers and dimers (Figure 1B, Supplementary Figure 2). The small amount of around 10% trimers and much fewer tetramers is likely caused by double-labeled proteins or nonspecific interactions. The significant number of monomers is likely caused by the necessary low concentrations in single-molecule fluorescence measurements. To minimize background signal, experiments must be performed at a minimum of nanomolar concentrations, which are frequently below the dissociation constant of protein oligomers [11,18]. Under these conditions, rebinding events are less likely to occur, biasing the observed distribution toward monomeric species. Consequently, the monomeric population of ArlJ detected here likely arises in part from these concentration limitations rather than reflecting the physiological oligomeric state of the protein. Moreover, the degree of labelling (DOL) of labelled ArlJ was around 66%, i.e. not all the proteins were labelled. To minimize the influence of this, the data were corrected for the DOL (see material and methods for more information), but a slight bias might still persist. Altogether our data show that ArlJ preferentially forms dimers rather than trimers. Including the previous finding that ArlI forms hexamers ([11]), the most plausible composition of the complex is a 2:6 assembly of ArlJ:ArlI.

**Figure 2.**
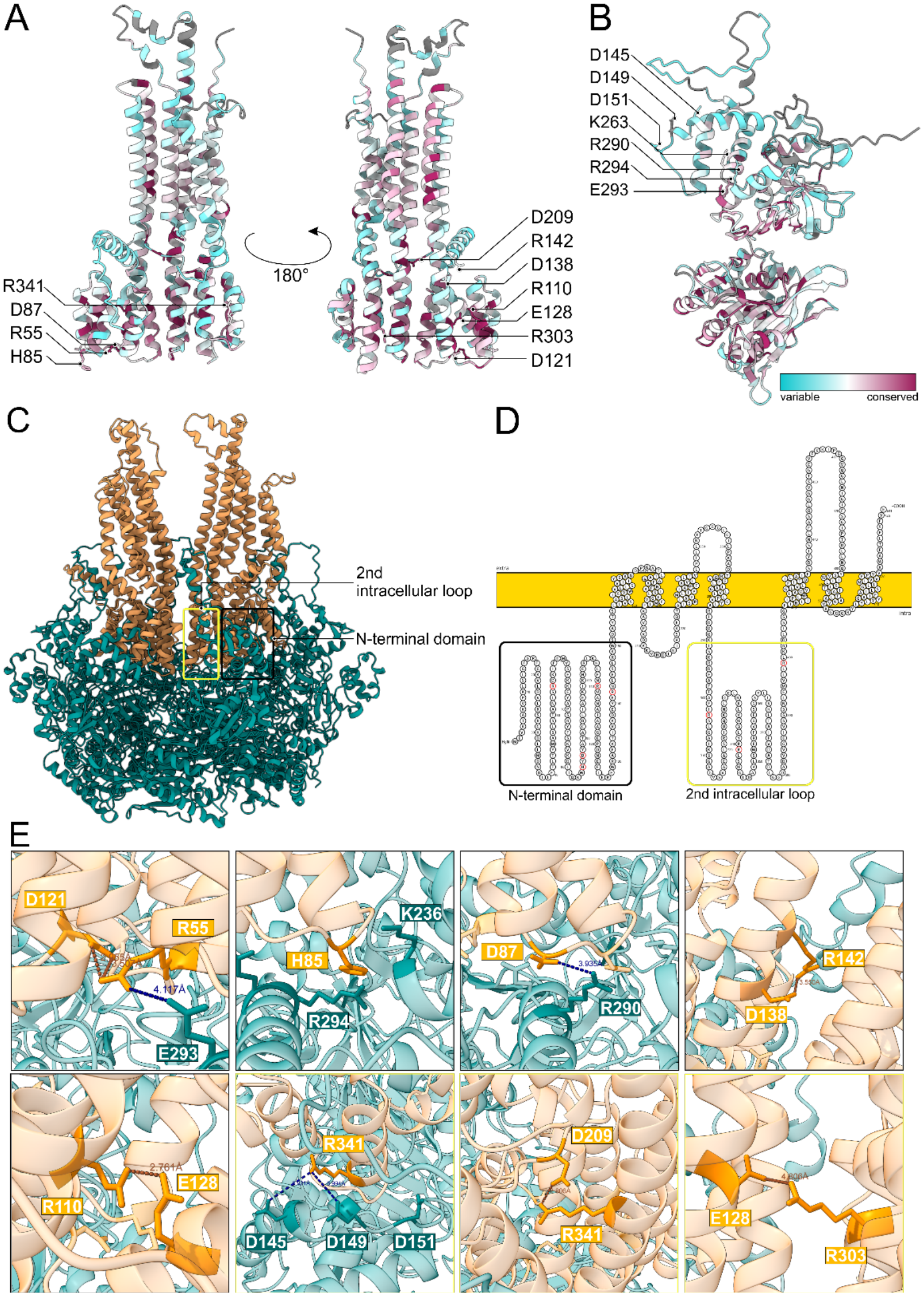
Conserved residues predicted to play a role in the ArlJ-ArlI interaction. **A.** Analysis of conserved residues in ArlJ. Residues of interest for further analysis are labelled. **B**. Analysis of conserved residues in ArlI. Labeled residues are potential interaction partners for oppositely charged and conserved residues in ArlJ. **C**. AlphaFold3 prediction of a dimeric ArlJ and a hexameric ArlI. The N-terminal domain of ArlJ is labelled in black, the 2^nd^ intracellular loop is labelled in yellow. **D**. Topology plot of ArlJ done with Protter [34]. The N-terminal domain is labelled in black, the 2^nd^ intracellular loop in yellow. Conserved residues of interest are marked in red. **E**. Zoomed-in view of the ArlI-ArlJ interaction sites. ArlJ is represented in orange, ArlI is represented in cyan. Conserved residues that are potential interaction partners for the ArlJ-ArlI complex formation are shown as atoms in the respective color. Distances between atoms are measured and shown in dark orange for intramolecular interactions (ArlJ-ArlJ) and intermolecular interactions (ArlJ-ArlI). Black boxes indicate location in the N-terminal domain, yellow boxes indicate localization in the 2^nd^ intracellular loop.

To corroborate our interpretation, we performed size-exclusion chromatography of ArlJ, which likewise resolved a single dimeric species as well, with no indication of a monomeric form (Supplementary Figure 1). Taken together, these complementary approaches support a dimeric arrangement of ArlJ distribution (Figure 1 B).

The addition of ATP did not significantly affect the oligomerization of ArlJ in complex with ArlI under the conditions tested, suggesting that nucleotide binding does not drive a change in the platform protein’s assembly state. These findings support a model in which ArlJ forms a dimeric membrane platform that associates with the hexameric ArlI complex (Figure 1 B, 5).

### Conserved residues in ArlJ are important for its interaction with ArlI *in vitro*

As structural data on Type IV pilus platform proteins remain limited, AlphaFold3 was used to investigate ArlJ–ArlI protein–protein interactions. Multiple sequence alignments of euryarchaeal ArlJ and ArlI were generated, and the predicted AlphaFold3 structures of the proteins of *Archaeoglobus fulgidus* were then coloured according to the conservation of individual residues (Figure 2 A, B). This analysis revealed moderate conservation of the long, protein-spanning helices of ArlJ, which are predicted to lie in the inner part of the oligomer. By contrast, the cytosolic loops of the protein are well conserved, and these loops are the regions most relevant for the ArlJ–ArlI interaction (Figure 2 A). In ArlI, the C-terminal Walker-A and Walker-B motifs are conserved, whereas the N-terminal domain displays a more variable region, consistent with previous analyses [9] (Figure 2 B).

The AlphaFold3 prediction further indicated that the candidate residues cluster in two distinct regions of ArlJ: the N-terminal domain and the second intracellular loop (Figure 2 C, D). R110, R142, R303 and R399 are positioned in relatively proximal cytosolic loops and are surrounded by nearby, oppositely charged residues, suggesting potential intra-protein interactions (Figure 2 E). In contrast, R55, H85 and D87 are located in more distal regions, making them more likely to participate directly in the interaction with ArlI (Figure 2 D, E). The AlphaFold3 model also identified conserved interaction partners within ArlI; notably, ArlJ R55 is found near the highly conserved D121 of a neighbouring helix. In addition, R341 lies in a separate loop and is surrounded by multiple negatively charged residues that form an acidic patch in the three-helix bundle of the ArlI NTD, although these residues are less conserved (Figure 2 D, E).

To test the role of these conserved residues in ArlJ, we set up a co-localization study. Co-expression plasmids of *A. fulgidus* ArlI and ArlJ were designed, and single-point mutations of the conserved residues were introduced (Supplementary Table 2). Following expression in *E. coli*, membrane isolation and pull-down of ArlJ via its His-tag, the co-localization of ArlI was assessed by SDS-PAGE (Figure 3 A). All amino acid exchanges reduced the amount of ArlI associated with ArlJ. Mutants predicted to engage in intramolecular interactions within ArlJ (R142, R303, R399) showed only minor effects, whereas point mutations predicted to lie in the more distal part of ArlJ (R55D, H85S, D87R, R341D) had markedly stronger effects. The largest decrease in ArlI relative to ArlJ was observed for R341D (Figure 3 B).

**Figure 3.**
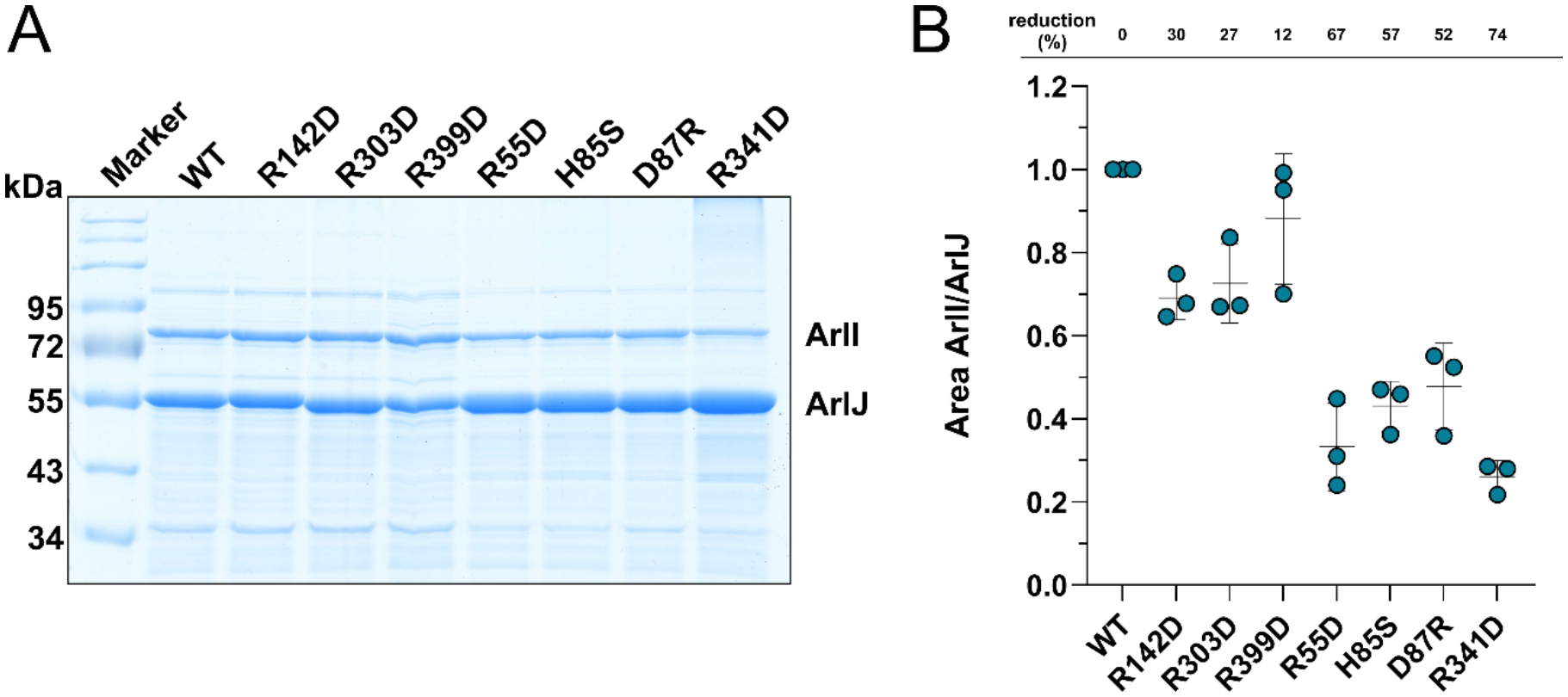
Co-localization studies with ArlJ mutants. **A.** Representative SDS-PAGE analysis of co-expressed and purified ArlJ-ArlI complex with single point mutations in ArlJ. **B**. Calculation of the ArlI/ArlJ ratio. Three individual biological replicates with the mean and standard deviation are shown.

### Conserved residues in ArlJ are important for swimming motility *in vivo*

The interaction studies of single-point ArlJ mutants described above indicated a clear effect on the ArlJ–ArlI interaction. Because these residues are conserved *ferax vocanii* was selected as the euryarchaeal model organism in which to investigate their contribution to swimming motility. The conserved residues identified in *A. fulgidus* had corresponding structural residues in *H. volcanii* ArlJ when the AlphaFold3 predictions were compared and structurally aligned (Figure 4 A).

**Figure 4.**
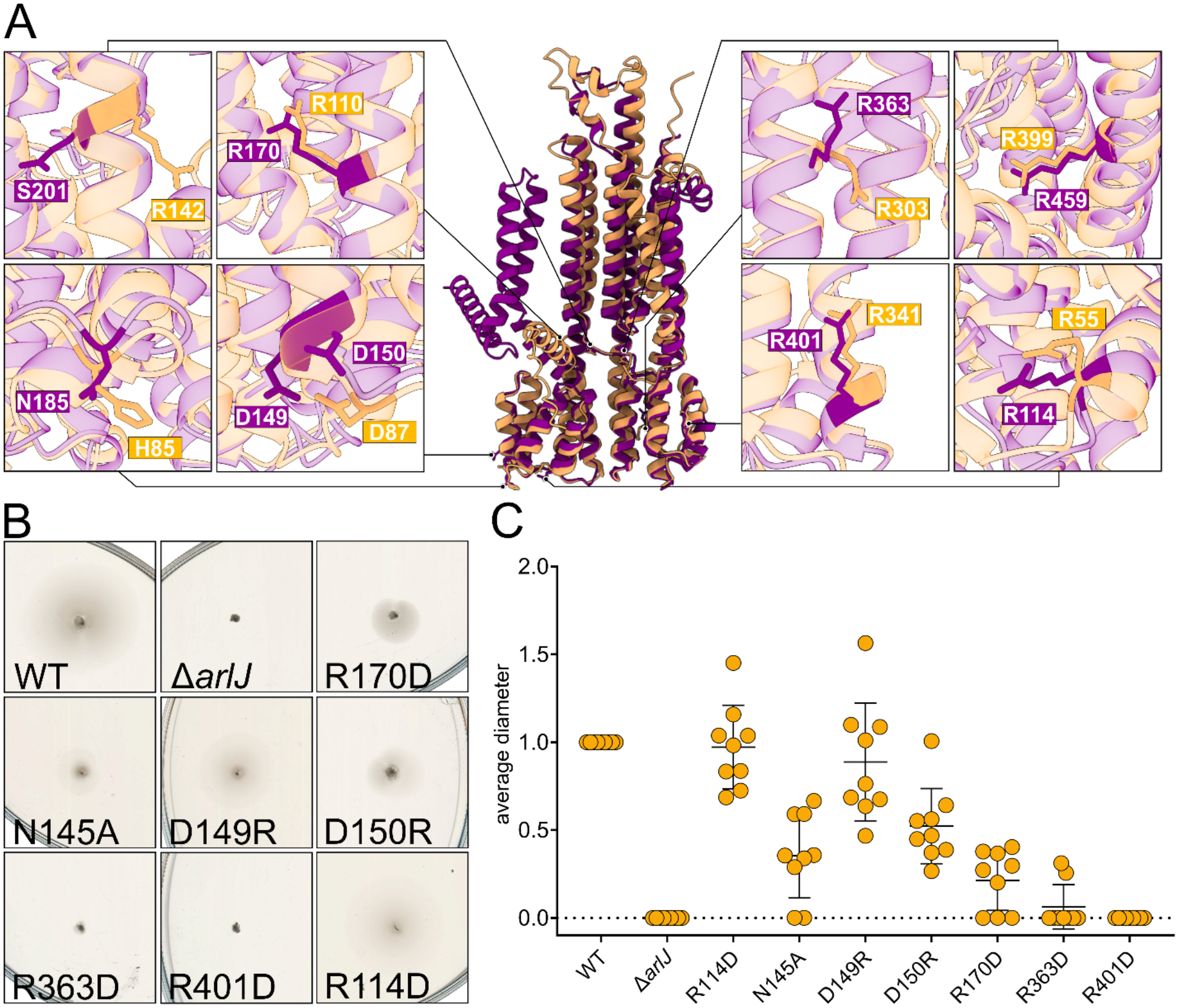
Conserved residues predicted to play a role in the ArlJ-ArlI interaction in *Haloferax volcanii*. **A.** Structural comparison of ArlJ of *H. volcanii* (purple) and *A. fulgidus* (orange). Residues of interest and their structural localisation are shown in detail in the close-up view. **B**. Motility analysis of single point mutations in *H. volcanii arlJ*.**C**. Calculated motility reduction in *arlJ* mutants in *H. volcanii*.

AfR303, AfR342, AfR399, AfR55 and AfR110 each had an explicit structural homolog in *H. volcanii* (Figure 4 A). For AfH85, the corresponding residue was N145, and for AfR142 it was S201. AfD87 had no direct charged structural counterpart but was flanked by two proximal aspartate residues. Single-point mutations of the corresponding residues were introduced into the genomic copy of *arlJ* in *H. volcanii* (Supplementary Table 1). The H26 wild-type strain exhibited motility consistent with previous reports [19], whereas the *arlJ* deletion strain was non-motile. (Figure 4 B). Among the point mutants, ArlJ R114D and D149R produced motility halos comparable to the wild type, suggesting that these residues are dispensable for archaellum rotation in *H. volcanii*. In contrast, the N145A, D150R, R170D, R363D, and R401D mutants showed a substantial reduction in motility, in line with the *in vitro* interaction data and reinforcing the functional importance of these conserved residues (Figure 4 B, C).

## Discussion

The rotation of the archaellum motor requires tight coupling between the cytoplasmic ATPase ArlI and the membrane platform protein ArlJ. Our data show that ArlJ forms a dimer that binds the oligomeric ArlI complex with a 2:6 stoichiometry (Figure 1 A) [11]. While functional information on T4P platform proteins remains sparse, the combination of conservation analysis, AlphaFold3-based structural predictions, heterologous co-purification and *in vivo* motility assays in *H. volcanii* allowed us to dissect the functional architecture of ArlJ and propose a mechanistic model for how ArlI-driven ATP hydrolysis is transduced into torque.

The cytoplasmic regions of ArlJ can be divided into two structurally and functionally distinct classes based on our mutation analysis: The first class (R110, R142, R303, R399) encompasses residues embedded in the proximal cytosolic loop, each of which is surrounded by oppositely charged residues consistent with intramolecular salt bridge formation. Their mutagenesis led to only a modest reduction in ArlI co-localization, yet mutation of structurally homologous residues in *H. volcanii* R170 and R363 caused severe motility defects. This dissociation between ArlI recruitment and motility is mechanistically informative: It argues that these residues maintain the structural integrity and geometry of the intracellular loop rather than directly contacting ArlI. A structurally perturbed loop can still recruit ArlI but fails to transmit force, consistent with a co-rotation model in which ArlJ loop rigidity is essential for torque transfer. Notably, hyper-motile mutations reported in close proximity to R363 in *H. volcanii* reinforce this view that subtle steric or electrostatic changes at this position can tune coupling efficiency in either direction, pointing to a mechanistically optimised, finely balanced interface [20].

The second class (R55, H85, D87 in the N-terminal domain and R341 in the 2^nd^ intracellular loop) occupies the distal tip of the two pseudo-symmetric loop elements and constitutes the direct ArlI interface. R341 stands out as the most critical single residue identified, as its substitution produced the largest reduction in colocalisation and abolished motility in *H. volcanii*. The predicted interaction partners of R341 are D145, D149 and D150 in ArlI and map directly to the three-helix bundle of the N-terminal ArlI domain, a region already implicated in membrane association [9]. Crucially, *Sulfolobus acidocaldarius* mutations in precisely M69, I72 and F76, which are structurally in similar positions to the *A. fulgidus* ArlI are equally deleterious, providing bidirectional mutational support for this salt-bridge contact as the primary load-bearing interface [9]. H85 and D87 in the N-terminal domain tip interact with an overlapping but distinct face of the three-helix bundle. H85 is particularly interesting: Histidine’s pH-sensitive ionisation state makes it a plausible state-dependent contact, capable of switching between cation-pi and hydrogen-bonding interactions, depending on the nucleotide state of ArlI, rather than forming a constitutive salt bridge [21].

Integrating this information within the known nucleo-tide-dependent conformational changes of the hexameric ArlI [9], we propose the following model: In the ATP-bound, closed conformation of ArlI, the three-helix bundle of the N-terminal domain approaches the second intracellular loop of ArlJ, engaging R341 against the acidic stripe of ArlI. This generates a tangential force component with a moment arm set by the radial distance of the R341 contact from the rotational axis. Simultaneously, H85 transitions to a more stable interaction with the same helix, contributing to force coupling. In the ADP/apo open state, the three-helix bundle withdraws and the N-terminal domain of the adjacent ArlJ protomer reengages via R55 and D87, stabilising the incoming ArlI subunit for the next power stroke. R55 may act bifunctionally, as its proximity to an internal ArlJ aspartate (D121) suggests an intramolecular ground-state role that primes the loop geometry before intermolecular engagement with ArlI. The overlapping contact phases between adjacent protomers smooth torque delivery and enable near-continuous rotation. This model is supported by the bleaching step analysis, which shows a 2:6 stoichiometry of ArlJ:ArlI independent of the nucleotide state of ArlI.

Altogether, our study puts forward a detailed mechanistic model of how force is transmitted from ATP hydrolysis in ArlI via conformational changes in ArlI (via power strokes) to rotation of ArlJ (Figure 5).

**Figure 5.**
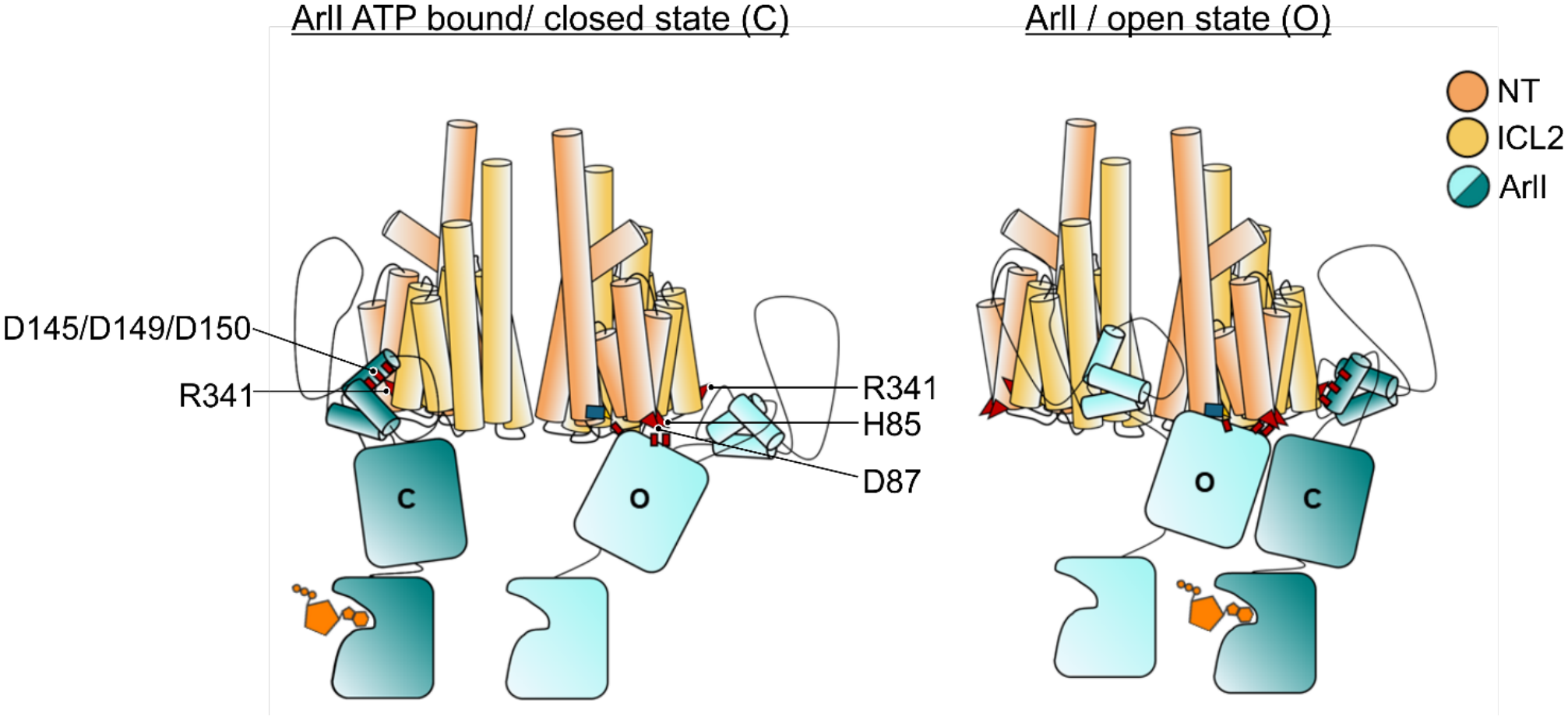
A model for the interaction of ArlI and ArlJ. Schematic representation of a dimeric ArlJ (orange) with ArlI (teal). In the ATP-bound state (closed conformation) ArlI binds to R341, providing a load-bearing coupling. In state 2, the same ArlI monomer is in an open state (ADP-bound or apo), in which the interaction occurs within the core area of the ArlI NTD, serving as a pre-recruitment/coordination state. The rotational force is created by tilting of the ArlI NTD throughout the process, generating a tangential force along the interaction axis with R341.

## Material and methods

### Strains

*H. volcanii* strains were grown in YPC medium rich medium (144 g l^−1^ NaCl, 21 g l^−1^ MgSO_4_*7H_2_O, 18 g l^−1^ MgCl_2_*6H_2_O, 4.2 g l^−1^ KCl, 12 mM Tris-HCl, 5 g l^−1^ yeast extract, 1 g l^−1^ peptone and 1 g l^−1^ Bacto™ Casamino acids or selective Hv-Ca medium.) or in selective Casamino acids medium (Hv-Ca; 144 g l^−1^ NaCl, 21 g l^−1^ MgSO_4_*7H_2_O, 18 g l^−1^ MgCl_2_*6H_2_O, 4.2 g l^−1^ KCl, 12 mM Tris-HCl 5 g l^−1^ Bacto™ Casamino Acids), both adjusted to pH 7.2 with KOH. When required, Hv-Ca medium was supplemented with 0.45 mM uracil [22]. Liquid cultures under 5 ml were grown in 15 ml tubes under constant rotation at 45 °C. Larger cultures were grown in baffled flasks at 120 rpm and 45 °C. Solid media plates were prepared as previously described [22] and incubated in sealed plastic containers at 45 °C to prevent evaporation. All strains used in this study are listed in Supplementary Table 1.

### Plasmids

Plasmids were cloned by restriction enzyme-based cloning or by in vivo ligation. All enzymes for DNA amplification (Phusion polymerase) and plasmid construction (restriction enzymes, T4 DNA ligase and Quick CIP) were obtained from New England Biolabs (NEB) and used according to the manufacturer’s protocol. Plasmids for co-expression of ArlJ and ArlI were cloned by site-directed mutagenesis.

To introduce single-point mutations of *arlJ* into *H. volcanii*, the existing knockout plasmid (pSVA5667) was used as a backbone, as it already contained the 500 bp homologous regions flanking *arlJ. ArlJ* was amplified from *H. volcanii* genomic DNA and cloned back between these flanking regions by Gibson Assembly, generating a plasmid for chromosomal knock-in of *arlJ*. Point mutations were subsequently introduced into this plasmid by mutagenesis PCR, yielding a series of knock-in plasmids each carrying a single amino acid exchange (Supplementary Table 2). Before transformation in *H. volcanii*, all plasmids were passed through a *dam*^*-*^ *E. coli* strain to obtain unmethylated DNA.

All plasmids are listed in Supplementary Table 2. Primers used for plasmid cloning are listed in Supplementary Table 3.

### Purification of ArlJ

*Escherichia coli* Rosetta (Novagen) cells transformed with pSVA13332 were used for heterologous expression of ArlJ. Cultures were grown in Terrific Broth (TB) at 37 °C with shaking at 120 rpm to an OD_600_ 0.5-0.6. Protein expression was induced with 500 µM IPTG for 4 hours under the same conditions. Cells were harvested by centrifugation and resuspended in resuspension Buffer (50 mM Hepes pH 7.0, 150 mM NaCl). Cell lysis was performed using a Microfluidizer (LM10 Microfluidizer, Microfluidics), followed by centrifugation at 27,200 x g for 15 min at 4°C. The supernatant was used for ultracentrifugation at 208,000 x g for 90 min at 4°C to isolate the membrane fraction. Membranes were resuspended in resuspension buffer and protein concentration was determined prior to solubilisation at 20-25 mg ml^−1^ protein in solubilization buffer (50 mM Hepes pH 7.0, 150 mM NaCl, 10 % glycerol, 1.5 % DDM) for 1 hour at 4°C. The solubilized sample was subsequently incubated at 65°C for 10 min to precipitate *E. coli* proteins, followed by ultracentrifugation at 208,000 x g for 45 min.

The clear supernatant was incubated with 1 ml of equilibrated NiNTA resin (cOmplete™ His-Tag Purification Resin, Roche) for 1 hour at 4°C. The resin was loaded onto a gravity column and the flow-through was collected. The column was washed with 20 column volumes (CV) wash buffer (50 mM HepespH 7.0, 150 mM NaCl, 10% glycerol, 0.03% DDM, 20 mM Imidazole). Bound protein was eluted with 7 CV elution buffer (50 mM Hepes pH 7, 150 mM NaCl, 10% glycerol, 0.03% DDM, 200 mM Imidazole). Eluted protein was concentrated to 500 µl in a 30 kDa Amicon and loaded to a Superdex 200 Increase 30/100 GL (Cytiva) size exclusion column, equilibrated in SEC buffer (50 mM Hepes pH 7, 150 mM NaCl, 10% glycerol, 0.03% DDM). Fractions containing ArlJ were pooled, concentrated in a 30 kDa molecular weight cut-off concentrator (Amicon, Merck Millipore Ltd.), snap frozen in liquid nitrogen and stored at -80°C until further usage.

### Purification of ArlI

For heterologous expression of *A. fulgidus* ArlI, *Escherichia coli* Rosetta (Novagen) cells transformed with pSVA13334 were used. Cultures were grown in Lysogeny Broth (LB) at 37 °C with shaking at 120 rpm to an OD_600_ 0.5-0.6. Protein expression was induced with 300 µM IPTG for 4 hours under the same conditions. In addition, 50 µM biotin was added to the cultures to enable biotinylation of the Avi-Tag. Cells were harvested by centrifugation and resuspended in resuspension Buffer (50 mM Hepes pH 8.0, 150 mM NaCl, 1 mM MgCl_2_). Cell lysis was performed using a french press (French® Pressure Cell, Glenn Mills) at 1000 psi. Lysed cells were centrifuged for 45 min at 27,200 x g at 4 °C. The supernatant was incubated with 1 ml of equilibrated NiNTA resin (cOmplete®™ His-Tag Purification Resin, Roche) for 1 hour at 4°C. The resin was loaded onto a gravity column and the flow through was collected. The column was washed with 20 (CV) wash buffer (50 mM Hepes pH 8.0, 150 mM NaCl, 1 mM MgCl2, 20 mM Imidazole). Bound protein was eluted with 7 CV elution buffer (50 mM Hepes pH 8.0, 150 mM NaCl, 1 mM MgCl_2_, 200 mM Imidazole). Eluted protein was concentrated to 500 µl in a 30 kDA molecular weight cut-off concentrator and loaded to a Superose6 Increase 30/100 GL (Cytiva) size exclusion column equilibrated in SEC buffer (50 mM Hepes pH 8, 150 mM NaCl, 1 mM MgCl_2_). Fractions containing ArlI were pooled, concentrated in a 30 kDA molecular weight cutoff concentrator (Amicon, Merck Millipore Ltd.), snap frozen in liquid nitrogen and stored at -80°C until further usage.

For heterologous expression of *P. furiosus* ArlI, *Escherichia coli* Rosetta (Novagen) cells transformed with pSVA13347 were used to inoculate Lysogeny Broth (LB) supplemented with 10 g l^−1^ Lactose and 50 µM biotin. Cultures were inoculated to an initial OD_600_ of 0.05 and incubated at 37 °C with shaking at 120 rpm for 16 hours. Protein purification was directly continued as described for *A. fulgidus* ArlI with the exception that purification was performed at 30 °C. For size exclusion chromatography, a Superdex 200 Increase 30/100 GL (Cytiva) was used. Fractions containing *P. furiosus* ArlI were pooled, concentrated in a 30 kDA molecular weight cut-off concentrator (Amicon, Merck Millipore Ltd.), and stored at 37°C with 25 µg mL^−1^ kanamycin.

### Labelling of ArlJ

*A. fulgidus* ArlJ^S388C^ was labelled with Alexa Fluor 647 C5 maleimide (ThermoFisher Scientific). Prior to labelling, the ArlJ cysteine was reduced with 10 mM TCEP (tris(2-carboxyethyl)phosphine) for 20 min on ice. TCEP was subsequently removed by five washing steps with ArlJ buffer (50 mM HEPES, 150 mM NaCl, 10% (m/v) glycerol, 0.009% (m/v) DDM, pH 7.0) using centrifugal filters (Amicon® Ultra, Merck Millipore, 80 kDa molecular weight cut-off). For the labelling reaction, a 1.5-fold molar excess of the fluorescent dye was added and the sample was incubated on a nutation plate overnight at 4 °C. Unbound dye was removed using a spin column (PD-25 MiniTrap, GE Healthcare) equilibrated in ArlJ buffer.

The degree of labelling (DOL) was determined from the protein and dye concentrations measured on a NanoDrop spectrophotometer (Thermo Scientific) and calculated according to eq. X. *f*_*c*_ is the correction factor of the dye, which corrects the absorbance at 280 nm, *A* is the absorbance, *c* the concentration, *ε* the extinction coefficient and *d* the pathlength.

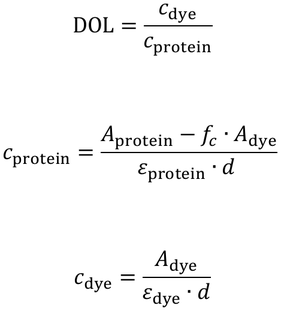

### Single-molecule TIRF experiment

The bleaching step experiments were conducted on a custom-built prism-type total internal reflection fluorescence (TIRF) microscope based on the setup described in Götz et al., 2018. The system was operated with flow chambers assembled from one quartz slide and one coverslip, with sample volume in between. The quartz glass was passivated with an 80:3 mixture of methoxy-PEG-silane (α-methoxy-ω-trimethoxysilylpropylurea PEG, 5 kDa, Rapp Polymere GmbH) and biotin-PEG-silane (3 kDa, Rapp Polymere GmbH), while the coverslip was fully passivated with methoxy-PEG-silane.

For the measurement, BSA (bovine serum albumin, 0.5mg/mL in HEPES-buffer, Carl Roth) was first flushed into the chamber and incubated for 1h to enhance the passivation. Subsequently, NeutrAvidin (0.25 mg/mL in HEPES-buffer, Thermo Fisher Scientific) was flushed in and incubated for 10 to 20 min to enable the immobilisation of the biotinylated protein. The chamber was then washed with ArlI buffer (50 mM HEPES, 150 mM NaCl, pH 8.0), followed by flushing in 200 nM or 200 pM biotinylated ArlI and incubation for 10 min. After washing with ArlJ buffer, 10 pM ArlJ was flushed in and incubated for another 10 min. Finally, unbound ArlJ was washed out with ArlJ buffer. For measurements containing ATP, 1 mM ATP and 5 mM MgCl_2_ were added to the ArlJ buffer, and 1 μM ArlJ was preincubated in this buffer for 1 h.

Excitation was performed by alternating laser excitation (ALEX) with either 200 ms red laser (637 nm, 5 mW, Coherent® OBISTM LX) [23], 50 ms dark time, 200 ms green laser (532 nm, 8 mW, Coherent® OBISTM LS) and 50 ms dark time, or with red laser illumination only (200 ms, 50 ms dark time) for 150 to 400 frames. Fluorescence was collected with a 100x oil immersion objective (Nikon) and recorded on an EMCCD camera (iXon Ultra 897, Andor) with 3x3 binning. Image sequences were stored as 16-bit TIFF stacks for subsequent analysis [23].

### Data analysis of bleaching step analysis

Traces showing bleaching steps were selected from intensity traces in IgorPro 6 with an in-house script. A trace was kept for analysis when its intensity drop led to a plateau lasting at least two frames (0.5 s), and when, in case of multiple bleaching steps, all plateaus had a similar height. The number of bleaching steps per trace was then determined by fitting the traces with AutoStepfinder [24]. Fits were generated by adjusting the parameters, and the fit best representing the data was chosen by eye. The fit results were processed with custom-written MATLAB scripts to obtain the occurrence of bleaching steps. The results were plotted as absolute and relative histograms. Because not 100% of the proteins are labelled with an appropriate number of fluorophores, the relative distribution was corrected for the degree of labelling (DOL) using a binomial distribution (eq. Y), where *n* is the oligomer size and *b* the number of observed bleaching steps. Since traces without any bleaching step, cannot be detected, the distribution was truncated at zero and renormalized (eq. Z). This analysis procedure follows the one described in [11].

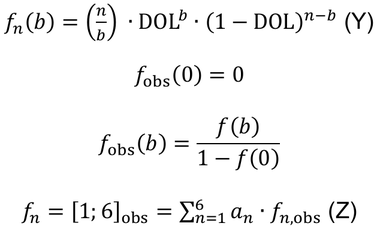

To account for the uncertainty of the DOL, which was estimated from several repeated NanoDrop (Thermo Scientific) measurements, the fitting was performed three times: with the DOL, and with the DOL plus or minus the confidence interval of 95%. The difference fits the length of the error bars. Each experiment was repeated three times, with 486 to 1000 traces.

### Bioinformatic analysis of conserved residues in ArlJ

To analyse conserved residues of ArlJ and ArlI, the amino acid sequence of both proteins was analysed using NCBI BlastP [25,26]. 1000 Target sequences were aligned in MEGA11 [27] using MUSCLE alignment [28], which was manually corrected by deleting mismatching sequences. This resulted in a n of 864 for ArlI and n=822 for ArlJ. For visualization, the structure of the AlphaFold3 prediction was coloured by the conservation in the multiple sequence alignment in ChimeraX [29] using the tool Colouring by Sequence conservation (entropy-based measure from AL2CO).

### Co-expression of ArlJ and ArlI

For heterologous co-expression of *A. fulgidus* ArlI and ArlJ, *E. coli* NiCo21 (New England Biolabs) cells transformed with the respective plasmid (Supplementary Table 2) were grown overnight in LB supplemented with 100 µg ml^−1^ ampicillin at 37 °C, 120 rpm. The overnight culture was used to inoculate 4x1 L TB to an OD_600_ of 0.05. Protein expression was induced with 300 µM IPTG at OD_600_ 0.5-0.7, the temperature was reduced to 30 °C and cultivation continued for 4 h. Cells were harvested by centrifugation at 6,250 x g at 4 °C, snap-frozen in liquid nitrogen and stored at - 80 °C until further usage.

The frozen pellet was resuspended in resuspension buffer (50 mM Hepes pH 7, 150 mM NaCl) supplemented with protease inhibitor (Pierce TM Protease Inhibitor Mini, Thermo Fisher Scientific ) and DNaseI (Roche) and lysed by two passages through the microfluidizer (LM 10 Microfluidizer, Microfluidics) at 1,500 bar. Cell debris was removed by centrifugation at 27,200 x g for 15 min at 4 °C. The supernatant was subjected to ultracentrifugation at 208,000 x g for 90 min at 4 °C to pellet the membrane fraction. The membrane pellet was resuspended in resuspension buffer and protein concentration determined by Bradford assay [30]. Membranes were solubilized for 90 min at 25 mg ml-1 total protein with 1.5% DDM (50 mM Hepes pH 7, 300 mM NaCl, 1 mM MgCl_2_, 10% glycerol, 1.5% DDM). After centrifugation at 208,000 x g for 45 min at 4 °C, the supernatant was incubated with equilibrated NiNTA resin (cOmplete™ His-Tag Purification Resin, Roche) for 90 min and loaded onto a gravity column. The column was washed with 20 CV wash buffer (50 mM Hepes pH 7, 300 mM NaCl, 1 mM MgCl_2_, 0.03 % DDM, 20 mM imidazole) and bound protein eluted with 10 CV elution buffer (50 mM Hepes pH 7, 300 mM NaCl, 1 mM MgCl_2_, 0.03 % DDM, 300 mM imidazole). The eluate was concentrated to 600 µl using a 100 kDA molecular weight cut-off concentrator (Amicon, Merck Millipore Ltd.). Concentrated protein was snap frozen in liquid nitrogen and stored at -80 °C until further usage.

### Co-localization studies

Co-expression of ArlJ and ArlI was analysed by SDS - Page and western blotting. Total protein concentrations were determined using a Bradford assay and adjusted to 600 µg ml-1. Samples were prepared with 5x SDS sample buffer (5:1 ratio) and 10 µl per sample (4.8 mg protein per lane) was loaded onto SDS-PAGE gels. Proteins were visualized by Q-stained SDS-APGE and detected in western blot using anti-his and anti-strep antibodies. Band intensities were quantified using ImageJ by integrating peak areas within each lane. The ratio of ArlJ/ArlI signal was calculated, normalised to the wild type and reported as mean +/-standard deviation from triplicate experiments.

### Mutant generation in *Haloferax volcanii*

Plasmids were transformed into *H. volcanii* Δ*arlJ* strain by PEG-600 mediated spheroplast transformation as previously described [31]. For transformation, *H. volcanii* Δ*arlJ* cells were grown to an OD_600_ 0.8 in 10 ml, pelleted (3,500 g, 10 min) and washed in buffered spheroplasting solution (1 M NaCl, 27 mM KCl, 50 mM Tris-HCl, 15% w/v sucrose, pH 8.5). Cells were resuspended in 600 µl of the same buffer and 200 µl aliquots used per transformation. Spheroplasts were formed by the addition of 50 mM EDTA (pH 8.0) for 10 min at room temperature. Unmethylated plasmid DNA (1-2 µg) was mixed with 15 µl unbuffered spheroplasting solution (1M NaCl, 27 mM KCl, 15% w/v sucrose, pH 7.5) and 5 µl 0.5 M EDTA, added to the cells and mixed by gentle inversion. After 5 min, 250 µl of 60% PEG600 (diluted in unbuffered spheroplasting solution) was added and the suspension was incubated for 30 min. Cells were then diluted in 1.5 ml spheroplasting dilution solution (23% saltwater, 15% w/v sucrose, 3.75 mM CaCl2) and pelleted (3,500 x g, 8 min). The pellet was overlaid with 1 ml regeneration solution (18% saltwater, 1x YPC, 15% w/v sucrose, 3 mM CaCl2) and incubated statically for 2 h at 45 °C, followed by 4 h under slow rotation at 45 °C. After final centrifugation (3,500 g, 8 min), cells were resuspended in 1 ml transformant dilution solution (18% saltwater, 15% w/v sucrose, 3 mM CaCl2) and 100 µl plated on selective Hv-Ca plates and incubated at 45 °C for 3 days.

Transformants were selected on Hv-Ca plates at 45 °C for three days. Selection is based on the *pyrE2* marker system, in which a plasmid carrying *pyrE2* and the homologous region of *arlJ* is transformed into the Δ*pyrE2*Δ*arlJ* strain, restoring uracil prototrophy upon chromosomal integration. To resolve the plasmid and leave only the mutated *arlJ* allele at the native chromosomal locus, transformants were serially passaged three times in 5 ml non-selective Hv-YPC liquid medium to promote intrachromosomal recombination and subsequent loss of the integrated plasmid. Depending on which side of the insertion the crossover occurs, this pop-out either restores the wild type or yields the desired mutant. In both cases, the cell loses pyrE2 and becomes uracil auxotroph, rendering it resistant to 5-FOA due to its inability to convert this toxic compound to the toxic analogue 5-fluorouracil. Pop-out clones were therefore selected on CA plates supplemented with 50 µg ml^−1^ 5-FOA and 0.09 mM uracil, plated at dilutions of 10^−1^,10^−2^ and 10^−3^. Single colonies were transferred to non-selective YPC plates and screened by colony PCR [22]. Correct chromosomal integration of each mutant was confirmed by sequencing and successful pop-out was additionally verified by the inability of clones to grow on uracil-free Hv-Ca plates.

### Motility assays

*Haloferax volcanii* was grown over night to an OD_600_ of 0.4-0.8. Semi-solid agar plates were prepared with CA-medium containing 0.3% agar and 50 ug/ml uracil. Cells diluted to an OD_600_ of 0.4 followed by spotting with a sterile toothpick piercing halfway through the agar. Plates were incubated for 5 days at 45 °C. Plates were scanned and analysed using Fiji ImageJ [32]. The average area of *H. volcanii* motility halos in relation to the H26 wild type was calculated for all replicates.

## Data availability

Data are provided in the main text, figures and supplementary data.

The single-molecule TIRF data for the bleaching step experiments are available from the authors upon request.

## Authors contributions

M.J and S.V.A conceived and designed the study with input from T.H. for single-molecule experiments. M.J. performed the in vitro experiments and protein purification for smTIRF, did data analysis and wrote the original manuscript. F.W. performed the in vitro and in vivo experiments and helped with data analysis. J.S., L.V. and E.S. established the smTIRF experimental setup and analyzed the data. R.K. constructed plasmids for the in vitro studies. T.H. supervised the smTIRF work. S.V.A. wrote the original manuscript. T.H. and S.V.A. acquired funding. All authors reviewed and edited the manuscript.

## Acknowledgements

We thank João Nuno de Sousa Machado for the ideas and initial groundwork that set the direction of this study. The archaellum shown in the graphical abstract was created in BioRender. Albers, S. (2027) https://BioRender.com/7qzqltf

