## Supplementary material for "More insights into the dynamic assembly and rotation of the archaellum motor machinery"

**Supplementary Table 1.** Strains used in this study.

| **Strain Name** | **Background** | **Genotype** | **Source/Reference** |
|---|---|---|---|
| ***Haloferax volcanii*** | | | |
| H26 | | Δ*pyrE2* | [1] |
| HTQ574 | H26 | Δ*pyrE2*Δ*arlJ* | This study |
| HTQ1102 | HTQ547 | Δ*pyrE2, arlJ*R114D | This study |
| HTQ1103 | HTQ547 | Δ*pyrE2, arlJ*N145A | This study |
| HTQ1104 | HTQ547 | Δ*pyrE2, arlJ*D149R | This study |
| HTQ1105 | HTQ547 | Δ*pyrE2, arlJD*150R | This study |
| HTQ1106 | HTQ547 | Δ*pyrE2, arlJ*R170D | This study |
| HTQ1107 | HTQ547 | Δ*pyrE2, arlJ*R363D | This study |
| HTQ1108 | HTQ547 | Δ*pyrE2, arlJ*R401D | This study |
| ***Escherichia coli*** | | | |
| NEB® 5-alpha Competent *E. coli* (High Efficiency) | - | *fhuA2::IS2 Δ(mmuP-mhpD)169 ΔphoA8 glnX44 ϕ80d[ΔlacZ58(M15)] rfbD1 gyrA96 luxS11 recA1 endA1 rphWT thiE1 hsdR17* | New England Biolabs |
| NiCo21 (DE3) | - | *can::CBD fhuA2 [lon] ompT gal (λ DE3) [dcm] arnA::CBD slyD::CBD glmS6Ala ΔhsdS λ DE3 = λ sBamHIo ΔEcoRI-B int::(lacI::PlacUV5::T7 gene1) i21 Δnin5* | New England Biolabs |
| Rosetta 2 (DE3) | - | $F^-$ *ompT hsdS*$_B$($r_B^-$ $m_B^-$) *gal dcm* (DE3) pRARE2 ($Cam^R$) | Novagen |
| *dam⁻/dcm⁻* | - | *ara-14 leuB6 fhuA31 lacY1 tsx78 glnV44 galK2 galT22 mcrA dcm-6 hisG4 rfbD1 R(zgb210::Tn10)* $Tet^S$ *endA1 rspL136* ($Str^R$)*dam13::Tn9* ($Cam^R$) *xylA-5 mtl-1 thi-1 mcrB1 hsdR2* | New England Biolabs |

**Supplementary Table 2. Plasmids used in this study.** Plasmids were cloned either by *in vivo* ligation (only primers listed) or restriction and ligation (primers and enzymes listed), as described in the methods section. AF, PF and Hvo stand for genes/proteins from *Archaeoglobus fulgidus*, *Pyrococcus furiosus* and *H. volcanii*, respectively.

| Plasmid | Function | Insert | Backbone | Primers used | Enzymes used | Source /Reference |
|---|---|---|---|---|---|---|
| pET-Duet-1 | Expression plasmid containing replicon ColE1 (pBR322) and two MCS (MCS1 and MCS2); $Amp^R$ | | | | | Novagen |
| **Plasmids for single molecule experiments** | | | | | | |
| pSVA5500 | Plasmid for expression of PF ArlI; $Amp^R$ | PF-6x*his-avi-arlI* | pET-Duet-1 | | | [2] |
| pSVA5456 | Plasmid for the expression of AF ArlJ $Amp^R$ | AF-6x*his-arlJ* (Insert was ordered from Invitrogen, see Suppl. Table 3) | pET-Duet-1 | 8404/8405 | BamHI/HindIII | This study |
| pSVA13332 | Plasmid for the expression of AF ArlJS388C used for single molecule experiments; $Amp^R$ | AF-6x*his-arlJS388C* | pSVA5456 | 12349/ 12350 | | This study |
| pSVA13334 | Plasmid for the expression of AF ArlJ | AF-6x*his-avi-arlI* | pSVA5500 | 12329/ 12330 12327/ 12328 | | This study |

| | | | | | | |
|---|---|---|---|---|---|---|
| | for single-molecule experiments; Amp$^R$ | | | | | |
| pSVA13347 | Plasmid for the expression of PF ArlIE24C for single-molecule experiments; Amp$^R$ | PF-6x*his*-*avi*-*arlI*E24C | pSVA5500 | 15000/ 15001 | | This study |
| **Plasmids for *in vitro* studies** | | | | | | |
| pSVA5461 | Plasmid for the co-expression of AF ArlJ-ArlI; Amp$^R$ | AF *his*-*arlJ*, *strep*-*arlI* (*arlJ* was already present in Backbone; *arlI* was ordered from Invitrogen, see Suppl. Table 3) | pSVA5456 | 8409/8408 | BamHI/AflIII | This study |
| pSVA13301 | Plasmid for the co-expression of AF ArlJR142D-ArlI; Amp$^R$ | AF *his*-*arlJR*142D, *strep*-*arlI* | pET-Duet-1 | 15007/ 15008 | | This study |
| pSVA13302 | Plasmid for the co-expression of AF ArlJR303D-ArlI; Amp$^R$ | AF *his*-*arlJR303*D, *strep*-*arlI* | pET-Duet-1 | 15009/ 15010 | | This study |
| pSVA13303 | Plasmid for the co-expression of AF ArlJR341-ArlI; Amp$^R$ | AF *his*-*arlJR341*D, *strep*-*arlI* | pET-Duet-1 | 15011/ 15012 | | This study |
| pSVA13304 | Plasmid for the co-expression | AF *his*-*arlJR399* | pET-Duet-1 | 15070/ 15071 | | This study |

| | | | | | | |
|---|---|---|---|---|---|---|
| | of AF ArlJR399D-ArlI; $Amp^R$ | D, *strep-arlI* | | | | |
| pSVA13351 | Plasmid for the co-expression of AF ArlJR55D-ArlI; $Amp^R$ | AF *his-arlJR55*D, *strep-arlI* | pET-Duet-1 | 15021/ 15022 | | This study |
| pSVA13356 | Plasmid for the co-expression of AF ArlJH85S-ArlI; $Amp^R$ | AF *his-arlJ*H85S, *strep-arlI* | pET-Duet-1 | 15032/ 15033 | | This study |
| pSVA13357 | Plasmid for the co-expression of AF ArlJRD87R-ArlI; $Amp^R$ | AF *his-arlJ*D87R, *strep-arlI* | pET-Duet-1 | 15034/ 15035 | | This study |
| **Plasmids for *in vivo* studies in *H. volcanii*** | | | | | | |
| pTA131 | Integrative plasmids for gene deletions in *H. volcanii* with a pyrE2 selection marker; $Amp^R$ | | | | | [3] |
| pSVA5667 | Knock-out plasmid for HVO *arlJ*; $Amp^R$ | HVO_1218 (*arlJ*) flanking regions | pTA131 | | KnpI/XbaI | This study |
| pSVA13371 | Plasmid for in-genome generation of single-point mutation in Hvo ArlJ; $Amp^R$ | HVO *arlJ* | pSVA5667 | 15050/ 15051 | | This study |
| pSVA13372 | Plasmid for in-genome generation of single-point | HVO *arlJ*R114D | pSVA13371 | 15054/ 15055 | | This study |

| | | | | | | |
|---|---|---|---|---|---|---|
| | mutation in Hvo ArlJR114D; $Amp^R$ | | | | | |
| pSVA13373 | Plasmid for in-genome generation of single-point mutation in Hvo ArlJN145S; $Amp^R$ | HVO *arlJ*N145S | pSVA13371 | 15056/ 15057 | | This study |
| pSVA13374 | Plasmid for in-genome generation of single-point mutation in Hvo ArlJN145A; $Amp^R$ | HVO *arlJ*N145A | pSVA13371 | 15057/ 15060 | | This study |
| pSVA13375 | Plasmid for in-genome generation of single-point mutation in Hvo ArlJD149R; $Amp^R$ | HVO *arlJ*D149R | pSVA13371 | 15062/ 15063 | | This study |
| pSVA13376 | Plasmid for in-genome generation of single-point mutation in Hvo ArlJD150R; $Amp^R$ | HVO *arlJ*D150R | pSVA13371 | 15064/ 15065 | | This study |
| pSVA13377 | Plasmid for in-genome generation of single-point mutation in Hvo ArlJR170D; $Amp^R$ | HVO *arlJ*R170D | pSVA13371 | 15066/ 15067 | | This study |

| | | | | | | |
|---|---|---|---|---|---|---|
| pSVA13378 | Plasmid for in-genome generation of single-point mutation in Hvo ArlJR363D; $Amp^R$ | HVO *arlJ*R363D | pSVA13371 | 15068/ 15069 | | This study |
| pSVA13379 | Plasmid for in-genome generation of single-point mutation in Hvo ArlJR401D; $Amp^R$ | HVO *arlJ*R401D | pSVA13371 | 15058/ 15059 | | This study |

**Supplementary Table 3.** Primers used in this study

| Primers | Sequence 5' → 3' | Description |
|---|---|---|
| 8404 | CGCCTGGGATCCGATGGAAGTTAAAACCGCAAAAG | Forward primer for amplification of AF *arlJ* with BamHI restriction site. Partner of 8405 to result in pSVA5456 |
| 8405 | GCGCGCAAGCTTTTATTCAACAAAGCTCGGC | Reverse primer for amplification of AF *arlI* with HindIII restriction site. Partner of 84084 to result in pSVA5456 |
| 8408 | TTGAATAAAAGCTTGCAGCATTAAGAAGGAGATATACCATGGGCTGGAGCCACCCGCAGTTCGAAAAAGGTAGCGAAGTTGATAAACTGGATTTTAGC<br><br>GCGGGCGCAAGCTTGCAGCATTAAGAAGGAGATATACCATGGGCTGGAGCCACCCGCAGTTCGAAAAAGGTAGCGAAGTTGATAAACTGGATTTTAGC | Forward primer for amplification of AF *arlI* with HindIII restriction site, ribosome binding site and Strep tag II sequence. Partner of 8409 to result in pSVA5461 |
| 8409 | GCAGGCGCCTTAAGTTACACTTTGATGGCCAGCAGAC | Reverse primer for amplification of AF *arlI* with AflII restriction site. Partner of 8408 to result in pSVA5461 |
| 10561 | CGGGGTACCGCGTCCGACATCGTCTCGATGATTC | Forward primer for amplification of genomic region upstream of Hvo *arlJ* with KpnI restriction site. Partner of 10562 to result in pSVA5667 |
| 10562 | TCTGTGGTCCGTCGTCAGTCATCTCGTTACACCCTCGCCATGTCGAACGG | Reverse primer for amplification of genomic region upstream of Hvo *arlJ* with complementary bases to downstream fragment (created with primers 10563,10564). Partner of 10561 to result in pSVA5667 |
| 10563 | ATGACTGACGACGGACCACAGAG | Forward primer for amplification of genomic region downstream of Hvo *arlJ*. Partner of 10563 to result in pSVA5667 |

| | | |
|---|---|---|
| 10564 | TGCTCTAGAGCGGAACAACAGCAGCGAGAAC | Reverse primer for amplification of genomic region of Hvo *arlJ* with XbaI restriction site. Partner of 10563 to result in pSVA5667 |
| 12329 | TTGAATGGCATGAAGGGTCCATGGGTAGCGAAGTTGATAAACTG | Forward primer for amplification of AF *arlI* with complementary bases for pSVA5500. Partner of 12330 to result in pSVA13334 |
| 12330 | CGCAAGCTTGTCGACCTGCAGTTACACTTTGATGGCCAGCAG | Revers primer for amplification of AF *arlI* with complementary bases for pSVA5500. Partner of 12329 to result in pSVA13334 |
| 12327 | GGACCCTTCATGCCATTCAATC | Reverse primer to linearise pSVA5500. Partner of 12327 to result in pSVA13334. |
| 12328 | CTGCAGGTCGACAAGCTTG | Forward primer to linearise pSVA5500. Partner of 12327 to result in pSVA13334. |
| 12349 | GTTAGCTGTAGCAACCTGGAAATGGTGCTG | Forward primer for site-directed mutagenesis of AF *arlJ* in pSVA5456, introducing S388C (AGC>TGT). Used with 12350 for whole plasmid amplification and *in vivo* ligation to result in pSVA13332 |
| 12350 | CAGGTTGCTACAGCTAACAATTTCACCCACAAC | Reverse primer for site-directed mutagenesis of AF *arlJ* in pSVA5456. Partner of 12349 to result in pSVA13332 |
| 15000 | CTAAGGAAATACGTTTGTGAGTTTAGGAGG | Forward primer for site-directed mutagenesis of Pf *arlI* in pSVA5500, introducing E24C (GAA>TGT). Used with 15001 for whole |

| | | |
|---|---|---|
| | | plasmid amplification and in vivo ligation to result in pSVA13347 |
| 15001 | <u>AACGTATTTCCTTAG</u>ATGAGGATTTCTTCTC | Reverse primer used for error prone PCR to result in pSVA13347 with <u>complementary bases</u> to pSVA5500/13347 |
| 15007 | CGCAAAGATGAATATGAA<u>GAT</u>AATCTGGAAAGCC | Forward primer for site-directed mutagenesis of *arlJ* in pSVA5461, introducing R142D (<u>CGC>GAT</u>). Used with 15008 for whole plasmid amplification and in vivo ligation to result in pSVA13301 |
| 15008 | <u>CATCTTTGCG</u>AATGGTTTTAAAGGCACGCC | Reverse primer for site-directed mutagenesis of arlJ in pSVA5461. Partner of 15007 to result in pSVA13301 |
| 15009 | GCATTTACCGCCTTTATT<u>GAT</u>AGCCTGGGTGC | Forward primer for site-directed mutagenesis fo *arlJ* in pSVA5461, introducing R303D (<u>CGC>GAT</u>). Used with 15010 for whole plasmid amplification and in vivo ligation to result in pSVA13302 |
| 15010 | <u>CGGTAAATGCT</u>TCATCACGTTTGCTGATTTTC | Reverse primer for site-directed mutagenesis of *arlJ* in pSVA5461. Partner of 15009 to result in pSVA13302 |
| 15011 | GGTTATGCAGCTGTATCGT<u>GAT</u>CTGAGCATGG | Forward primer for site-directed mutagenesis of *arlJ* in pSVA5461, introducing R341D (<u>CGC>GAT</u>). Used with 15012 for whole plasmid amplification |

| | | |
|---|---|---|
| | | and in vivo ligation to result in pSVA13303 |
| 15012 | CAGCTGCATAACCAGTTCACGCAGTTCACC | Reverse primer for site-directed mutagenesis of *arlJ* in pSVA5461. Partner of 15009 to result in pSVA13303 |
| 15021 | CTCAGATTTGCGGTGCTCAGG | Reverse primer for site-directed mutagenesis of *arlJ* in pSVA5461. Partner of 15022 to result in pSVA13351 |
| 15022 | CGCAAATCTGAGCGATGATAAAATCTTTG | Forward primer for site-directed mutagenesis of *arlJ* in pSVA5461, introducing R55D (CGT>GAT). Used with 15021 for whole-plasmid amplification and in vivo ligation to result in pSVA13351 |
| 15032 | CTGACCCAGAAATGGTCATATGATTATGC | Forward primer for site-directed mutagenesis of *arlJ* in pSVA5461, introducing H85S (CAT>TCA). Used with 15033 for whole-plasmid amplification and in vivo ligation to result in pSVA13356 |
| 15033 | CCATTTCTGGGTCAGATCTTTCACTTTG | Reverse primer for site-directed mutagenesis of *arlJ* in pSVA5461. Partner of 15032 to result in pSVA13356 |
| 15034 | ATAATGCCATTTCTGGGTCAGATCTTTCAC | Reverse primer for site-directed mutagenesis of *arlJ* in pSVA5461. Partner of 15035 to result in pSVA13304 |
| 15035 | CAGAAATGGCATTATCGTTATGCAACCG | Forward primer for site-directed mutagenesis of *arlJ* in pSVA5461, |

| | | |
|---|---|---|
| | | introducing D87R (GAT>CGT). Used with 15033 for whole-plasmid amplification and in vivo ligation to result in pSVA13357 |
| 15050 | GAGGGTGTAACGAGATGGCAGGGGAACAG | Forward primer for amplification of Hvo *arlJ* with complementary bases for pSVA5667 |
| 15051 | GGTCCGTCGTCAGTCATACACTAATCAG | Reverse primer for amplification of Hvo *arlJ* with complementary bases for pSVA5667 |
| 15054 | CGACGAACATCGACGACGTCGCCGTG | Forward primer for site-directed mutagenesis of Hvo *arlJ* in pSVA13371, introducing R114D (GCG>GAC). Used with 15055 for whole-plasmid amplification and *in vivo* ligation to result in pSVA13372 |
| 15055 | GTCGATGTTCGTCGTCGAGAGCACG | Reverse primer for site-directed mutagenesis of Hvo *arlJ* in pSVA13371. Partner of 15054 to result in pSVA13372 |
| 15056 | CTCGTCGACGCGTGGTCGCAGTCG | Forward primer for site-directed mutagenesis of Hvo *arlJ* in pSVA13371, introducing N145S (AAC>TCG). Used with 15057 for whole-plasmid amplification and *in vivo* ligation to result in pSVA13373 |
| 15057 | CCACGCGTCGACGAGCTGGACGATG | Reverse primer for site-directed mutagenesis of Hvo *arlJ* in pSVA13371. Partner of 15056 to result in pSVA13373 |
| 15058 | GCCCGGCTGTACACGGACCTTCGG | Forward primer for site-directed |

| | | |
|---|---|---|
| | | mutagenesis of Hvo *arlJ* in pSVA13371, introducing R401D (CGC>GAC). Used with 15059 for whole-plasmid amplification and *in vivo* ligation to result in pSVA13379 |
| 15059 | CGTGTACAGCCGGGCGATTTCACGC | Reverse primer for site-directed mutagenesis of Hvo *arlJ* in pSVA13371. Partner of 15058 to result in pSVA13379 |
| 15060 | CTCGTCGACGCGTGGGCCCAGTCGC | Forward primer for site-directed mutagenesis of Hvo *arlJ* in pSVA13371, introducing N145A (AAC>GCC). Used with 15057 for whole-plasmid amplification and *in vivo* ligation to result in pSVA13374 |
| 15062 | GTGGAACCAGTCGCTCCGCGACGCC | Forward primer for site-directed mutagenesis of Hvo *arlJ* in pSVA13371, introducing D149R (GAC>CGC). Used with 15063 for whole-plasmid amplification and *in vivo* ligation to result in pSVA13375 |
| 15063 | GAGCGACTGGTTCCACGCGTCGACG | Reverse primer for site-directed mutagenesis of Hvo *arlJ* in pSVA13371. Partner of 15062 to result in pSVA13375 |
| 15064 | GAACCAGTCGCTCGACCGCGCCTGTC | Forward primer for site-directed mutagenesis of Hvo *arlJ* in pSVA13371, introducing D150R (GAC>CGC). Used with 15065 for whole-plasmid amplification and *in vivo* ligation to result in pSVA13376 |

| | | |
|---|---|---|
| 15065 | GTCGAGCGACTGGTTCCACGCGTCG | Reverse primer for site-directed mutagenesis of Hvo *arlJ* in pSVA13371. Partner of 15064 to result in pSVA13376 |
| 15066 | GCCGACTTCCTCGACGACCTCGCG | Forward primer for site-directed mutagenesis of Hvo *arlJ* in pSVA13371, introducing R170D (CGC>GAC). Used with 15067 for whole-plasmid amplification and *in vivo* ligation to result in pSVA13377 |
| 15067 | GTCGAGGAAGTCGGCGAGCGGCTTC | Reverse primer for site-directed mutagenesis of Hvo *arlJ* in pSVA13371. Partner of 15066 to result in pSVA13377 |
| 15068 | GTACCCCGGCTTCATCGACGCGCTCG | Forward primer for site-directed mutagenesis of Hvo *arlJ* in pSVA13371, introducing R363D (CGC>GAC). Used with 15069 for whole-plasmid amplification and *in vivo* ligation to result in pSVA13378 |
| 15069 | ATGAAGCCGGGGTACTCCTCGTCGC | Reverse primer for site-directed mutagenesis of Hvo *arlJ* in pSVA13371. Partner of 15068 to result in pSVA13378. |
| 15070 | GGTGCTGCTGGATCTGAAACGCG | Forward primer for site-directed mutagenesis of *arlJ* in pSVA5461, introducing R399D (CGC>GAT). Used with 15071 for whole-plasmid amplification and in vivo ligation to result in pSVA13304 |
| 15071 | CAGCAGCACCATTTCCAGGTTGCTG | Reverse primer for site-directed |

| | | |
|---|---|---|
| | | mutagenesis of *arlJ* in pSVA5461. Partner of 15070 to result in pSVA13304 |

**Supplementary Table 4.** Gene sequences ordered from Invitrogen for heterologous expression in *E. coli*.

| Gene | Sequence |
|---|---|
| AF_1049 (*Archaeoglobus fulgidus arlI*) | GGATCCATGGGTAGCGAAGTTGATAAACTGGATTTTAGCCTGAAACCGAAAGGTCTGAAA<br>CAGGGTGTTGTTGCAAATTTTCCGTTTAAACCGCGTGTTCTGGAAGAAGATCACGATCAT<br>AGCAATCTGGCAAGCTGTGGTATCTATAAACTGCTGCCGGATCGTATGAAAAAAGAGGTG<br>GAAAAAAATCTGCATCTGCTGGAATATCTGCATATTCTGCCGCTGGATAAAATTGGCATT<br>CCGAAATTTGCACCGAAACTGGATCGTAAAAAACATGGTGATCTGGATCGTCCGAATCTG<br>ATTTATCCGGCAGATGAGCAGATTTATATCCATGTTTATCCGGACAAAAACGATATGCGC<br>AACTTTTATATCCCGATTGAACCGACACTGCTGACCGGTGTTGATGAACTGCTGAAAGAA<br>GTTGAAATTCGTCTGGTGGATTATATCACCGATATCGATTTTGACCCGGAAGATAACGAA<br>GAAAAGATCAAGATTCTGAAAGAGGCACTGCGCGAAGTTTGTGAAATTGTTGATAAAGAA<br>GAATCCTTTATGGATGATAGCCGTGGTGAAGTTAAACTGACCCCTAAATTTCGCTTTCCG<br>TTTCGTCTGCCGTTTTGGAAAAATGGTAAAAATGGCAGCCGTGCCGTTATGGGTAAACTG<br>AAAGTTACCAAAGAACAGTTCAAAGCCCTGGAATATGCACTGATTCGTGATAAAGTTGGT<br>CTGGGTGTGCTGGAACCGTATATTCGCGATAAATACATTGAGGATATTAGCTGTAGCGGT<br>CTGGGTCCGATTTTTGTGGAACACAAAATCTTTAAAGGCCTGAAAAGCGTGATTGAATTT<br>CGTGATGAAGAAACCCTGAACAAGTTTATTATCCGTCTGGCAGAACGTATTGGTAAACCG<br>GTTACCTATAAAGATCCGATTGTTGATGCAACCCTGCCGGATGGTAGCCGTATTAACATT<br>GTTTTTGGCAACGACATTAGCAAGAACGGCAGCAATTTTACCATCCGCAAATTTAACGAA<br>ACCCCGTTTAGCGTTCTGCAACTGATCGAATTTGGTAGCATCGATTATATGATTGCCGGT<br>TATCTGTGGCTGCTGATTAGCGAAGGTATGAGCGGTTTTATTTGTGGTGAAACCGCAAGC<br>GGTAAAACAACCCTGCTGAATGCAATTACCGCATTTATTCGTCCGGAAGCAAAAGTTGTT<br>AGCATTGAAGATACACCGGAACTGCAAGTGCCGCATAAAAACTGGACCCGTGAAGTTACC<br>CGTGGTAGCACCCGTGCCGGTGGTATTGGTGAAGGTGGTGGTAGTGATGTTACCATGTTT<br>GATTTACTGAAAGCAGCCCTGCGTCAGCGTCCGAACTATATTCTGGTTGGTGAAATTCGC |

| | |
|---|---|
| | GGTGTGGAAGGTAATATTGCATTTCAGGCAATGCAGACCGGTCA<br>TCCGGTTATGAGCACC<br>TTTCATGCAGCAACCGTTGAAAAACTGATTCAGCGTCTGACCAC<br>CGAACCGATTAGCGTT<br>CCGAAAACCTTTATTGATAACCTGAACTTCGTTGTGATTCAGAGC<br>GCAGTTCGTCGTCCG<br>GATGGCAAACTGGTTCGTCGTGTTCTGAGCGTTAGCGAAATTGT<br>GGGTTATAATCCGCAG<br>AAAGGTGGTGTGAGCTTTATTGAAGTTTTTCAGTGGGACCCTGT<br>TACCGATACACATGTT<br>TTTACCGGTTATGGTAGCAGCTACCTGCTGGAACAGAAAATTGCA<br>ACCCGTCTGGGTATT<br>CCGCCTAATAAGAAAAAGCTGATTTACGAAGAGGTTGAGAAACG<br>TGCCAAAATCCTGAAA<br>AAACTGCATGAACAGGGCGTGACCGATTTTTATGAATTTTTCAAA<br>AGCATCAGCAAAATC<br>TATAAAAGCGGTCTGCTGGCCATCAAAGTGTAAAAGCTT |
| AF_1048 (*Archaeoglobus fulgidus arlJ*) | GGATCCATGGAAGTTAAAACCGCAAAAGTTCCGAAACTGAGCTT<br>AGGTTTTGGTCGTTTT<br>GGTAAAGTTCTGGATGTTCTGCGTAGCCGTGGTAAAGATCTGCT<br>GATGGATAATGATCTG<br>CTGTTTACCCTGACCTATATGGCAAGCCTGAGCACCGCAAATCT<br>GAGCCGTGATAAAATC<br>TTTGAAATGGTGAGCGAGAAGAAAGAATATGCACCGAGCCGTTA<br>TTTCAGCAAAGTGAAA<br>GATCTGACCCAGAAATGGCATTATGATTATGCAACCGCATGTGAT<br>CTGATCGCCGAAAAA<br>ATCAAACATGAGCGTCTGAAAAAACTGTTTAATCGTATGGCAAAT<br>GCCATTGCAGCCGGT<br>GAACCGGATAATGAATTTCTGGAACGTGAATGGCGTGCCTTTAAA<br>ACCATTCGCAAAGAT<br>GAATATGAACGCAATCTGGAAAGCCTGCGTAAATGGTCAGATGC<br>ATATACCAGCCTGCTG<br>GTTAGCGCCAGCCTGATTAGCGTTGTTGTTCTGCTGAGCGTTAT<br>TATCTATAGTGCCGGT<br>GATCCGGGTGCAACCCTGACCGCAAGCGCCTTTGCAAATTTTG<br>CCATTAGCCTGTTTGGT<br>GTGTTCATGCTGTTTAAAGCCGTTCCGAAAGATAAAAAAGTGCAC<br>GACCTGAAGATCAAG<br>AGCAAAGAACAGACCGCAATTAGCCGTCTGGCACCGATGCTGAT<br>TCCGATTGCATTTTTT<br>GCAGTTCTGTTTCTGACCGTTCTGCCGACACTGCTGAATTTTAC<br>CGGTGGTGGTCTGGGT<br>ATTCTGCCGAAAGCAGATTATCGTGGTGTTGGTTTTCTGCTGGC<br>AGGTCTGGTTCTGCTG<br>CCGGTTGGTTATCTGGCACGTATTGATGATCAGAAAATCAGCAAA<br>CGTGATGAAGCATTT<br>ACCGCCTTTATTCGTAGCCTGGGTGCAATTAAAAGCGGTGCCGG<br>TGTTAGCGTTGCAGAA |

| | |
|---|---|
| | GCACTGAGCCGTATTGATCAAAAAAATCTGGGTGAACTGCGTGA<br>ACTGGTTATGCAGCTG<br>TATCGTCGTCTGAGCATGGGTTTAGATGCAAAACTGAGTTGGGA<br>ACGTTTTGTTGGTGAA<br>AGCGGTAGCTATCTGATTAGCAAACTGACCGCCATTTTTGTTGAT<br>GCAACCGATTTAGGT<br>GGTGATGCAGATGTTGTGGGTGAAATTGTTAGCAGCAGCAACCT<br>GGAAATGGTGCTGCTG<br>CGCCTGAAACGCGATCTGATTTCAAGCGGTTTTATCAACCTGATT<br>ATTCCGCTGCATCTG<br>GCAATGGTTGGTCTGGTGCTGTTTATTACCCAGATTCTGGCAATT<br>TTCAGCGACTATATT<br>AGCAACCTGTTTGCAAGCCAGCTTGGTGGTGTTAGCACCGGTG<br>AAGTTTTTAGCAAAATT<br>CCTGGTGGTATGCAGGGTCTGAATATTGGTATTTTTAGCGGTATT<br>CCGACAGACCTGCTG<br>AATCAGTATAGCGTTTGGATTATTCTGAGCCTGACCATTGCAAATA<br>CCCTGGCAGCAAAT<br>GTTGTTAAAGGTGGTGGCCGTTATCTGTATCTGTATTATGGTGCA<br>ATTTTTGCGATTCTG<br>AGTGGCCTGCTGATGCTGGTTGTTCCTCCGGTTGTTAAATGGGC<br>ATTTACCCTGCCGAGC<br>TTTGTTGAATAAAAGCTT |

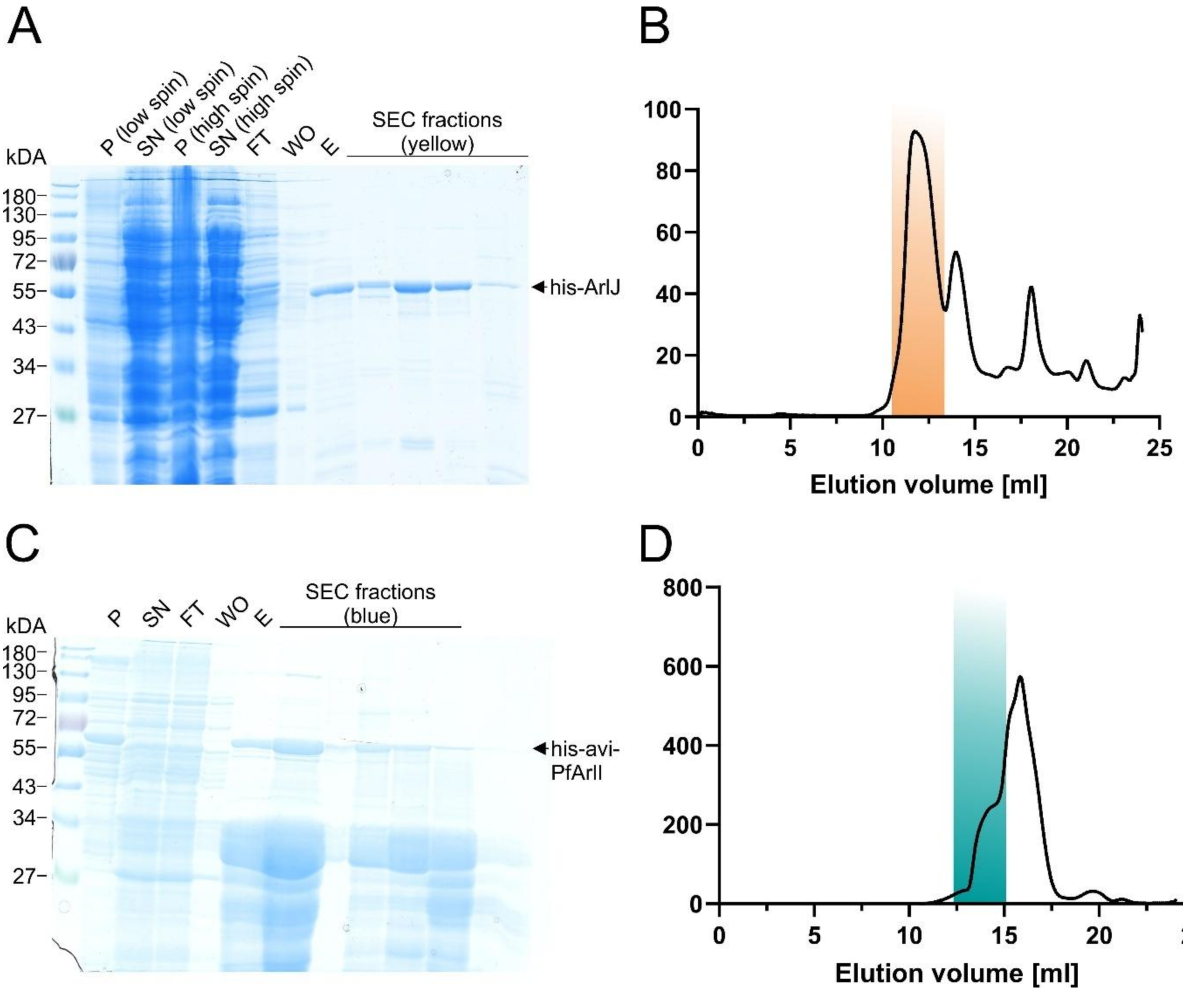


**Supplementary Figure 1**. **Purification of AF ArlJ (A+B) and PF ArlI (C+D).** (A) Coomassie-stained SDS-PAGE showing the AF ArlJ purification as described in the methods section. Lanes from left to right: pellet (P), Supernatant (SN), NiNTA column flow-through (FT), NiNTA column wash-out (WO), elution fraction of the NiNTA column and size exclusion fraction as indicated in yellow in (**B**). (**B**) Size-exclusion chromatogram of AF ArlJ purification. Yellow indicates the volume collected for the single-molecule measurements and shown on the SDS-PAGE (**A**). (**C**) Coomassie-stained SDS-PAGE showing the PF ArlI purification. From left to right: pellet (P), Supernatant (SN), NiNTA column flow-through (FT), NiNTA column wash-out (WO), elution fraction of the NiNTA column and size exclusion fraction as indicated in blue in (**D**). (**D**) Sie exclusion chromatogram of the PF ArlI purification. Blue indicates the collected volume used for the single-molecule measurements and shown on the SDS-PAGE (**A**).

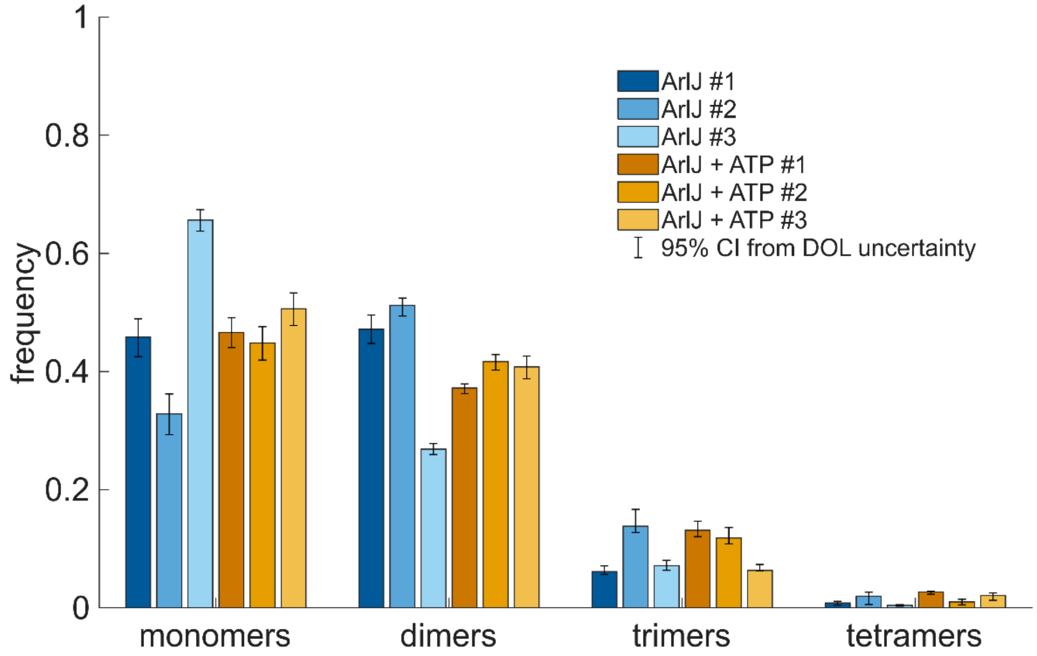


**Supplementary Figure 2.** Individual measurements underlying the mean frequencies shown in Figure 1B. Oligomer distributions of ArlJ for each independent experiment (n=3) in the absence (blue) and presence (orange) of ATP. The data was corrected for a DOL of 66 %. Measurements ArlJ #1, ArlJ #2 and ArlJ + ATP #1 were performed using 200 nM ArlI and 10 pM ArlJ and the measurements ArlJ #3, ArlJ + ATP #2 and ArlJ + ATP #3 were performed with 200 pM ArlI and 10 pM ArlJ.
